# Stable recombination hotspots in birds

**DOI:** 10.1101/023101

**Authors:** Sonal Singhal, Ellen M. Leffler, Keerthi Sannareddy, Isaac Turner, Oliver Venn, Daniel M. Hooper, Alva I. Strand, Qiye Li, Brian Raney, Christopher N. Balakrishnan, Simon C. Griffith, Gil McVean, Molly Przeworski

## Abstract

Although the DNA-binding protein PRDM9 plays a critical role in the specification of meiotic recombination hotspots in mice and apes, it appears to be absent from many vertebrate species, including birds. To learn about the determinants of fine-scale recombination rates and their evolution in natural populations lacking *PRDM9*, we inferred fine-scale recombination maps from population resequencing data for two bird species, the zebra finch *Taeniopygia guttata*, and the long-tailed finch, *Poephila acuticauda*, whose divergence is on par with that between human and chimpanzee. We find that both bird species have hotspots, and these are enriched near CpG islands and transcription start sites. In sharp contrast to what is seen in mice and apes, the hotspots are largely shared between the two species, with indirect evidence of conservation extending across bird species tens of millions of years diverged. These observations link the evolution of hotspots to their genetic architecture, suggesting that in the absence of PRDM9 binding specificity, accessibility of the genome to the cellular recombination machinery, particularly around functional genomic elements, both enables increased recombination and constrains its evolution.

## One-Sentence Summary

We show the fine-scale recombination landscape is stable across tens of millions of years in birds, in sharp contrast to what is seen in primates and mice.

## Main Text

Meiotic recombination is a ubiquitous and fundamental genetic process that shapes variation in populations, yet our understanding of its underlying mechanisms is based only on a handful of model organisms, scattered throughout the tree of life. One pattern shared among most sexually reproducing species is that meiotic recombination tends to occur in short segments of 100s to 1000s of base pairs, termed “recombination hotspots” (*1*). In apes and mice, the location of hotspots is largely determined by PRDM9, a zinc-finger protein that binds to specific motifs in the genome during meiotic prophase and generates H3K4me3 marks, eventually leading to double-strand breaks (DSBs) and both crossover and non-crossover resolutions (*2-5*). The zinc-finger domain of *PRDM9* evolves quickly, with evidence of positive selection on residues in contact with DNA (*2, 6*), and as a result, there is rapid turnover of hotspot locations across species, subspecies, and populations (*7-9*).

Whereas *PRDM9* plays a pivotal role in controlling recombination localization in mice and apes, many species lacking *PRDM9* nonetheless have hotspots (*6*). An artificial example is provided by mice knockouts for PRDM9. Although sterile, they make similar numbers of DSBs as wild-type mice, and their recombination hotspots appear to default to residual H3K4me3 mark locations, notably at promoters (*10*). A natural but puzzling example is provided by canids, who carry premature stop codons in *PRDM9,* yet are able to recombine and remain fertile (*11*). Like mouse *PRDM9* knockouts, in canids and in other species without *PRDM9* such as *Saccharomyces cerevisae* and *Arabidopsis thaliana*, hotspots tend to occur at promoters or other regions with promoter-like features (*11-13*). In some taxa without *PRDM9*, notably *Drosophila* species (*14, 15*), honeybees (*16*), and *Caenorhabditis elegans (17)*, short, intense recombination hotspots appear to be absent.

To further explore how the absence of PRDM9 shapes the fine-scale recombination landscape and impacts its evolution, we turn to birds, because they have compact and largely syntenic genomes that facilitate comparative study (*18*) and because an analysis of the chicken genome suggested that birds may not have *PRDM9* (*6*). We first confirmed the absence of *PRDM9* in birds by querying the genomes of 48 species of birds, three species of crocodilians, two species of turtles, and one species of lizard for *PRDM9* (*19, 20*). We found that only turtles contain putative orthologs with all three *PRDM9* domains (Fig. S1). We also found no expression of any *PRDM9*-like transcripts in RNAseq data from testes tissue of the zebra finch (*Taeniopygia guttata*). Given the likely absence of *PRDM9* in birds, we ask: is recombination nonetheless concentrated in hotspots in these species? If so, how quickly do the hotspots evolve? Where does recombination tend to occur in the genome? To address these questions, we generated whole-genome resequencing data for wild populations from two bird species and inferred fine-scale genetic maps from patterns of linkage disequilibrium. While this approach only generates a historical, sex-averaged map of recombination (*21*), other methods such as sperm typing, double strand break sequencing, or analyses of pedigrees are impractical for most non-model species (*22*).

### Inferring fine-scale recombination maps

We generated whole genome resequencing data from wild populations of three species of finch in the family *Estrildidae*: zebra finch (*Taeniopygia guttata*; n=19 wild, unrelated birds and n=5 from a domesticated, nuclear family), long-tailed finch (*Poephila acuticauda*; n=20, including 10 of each of two, similar subspecies with average autosomal F_ST_ = 0.039), and double-barred finch (*Taeniopygia bichenovii*; n=1) (Fig. 1, Table S1; (*23*)). Despite extensive incomplete lineage sorting between the species, there is no evidence for gene flow amongst them (Fig. S2). Moreover, nucleotide divergence among the three finch species is similar to that of human, chimpanzee and gorilla, providing a well-matched comparison to apes (*8, 9*) (Fig. 1).

**Figure 1:**
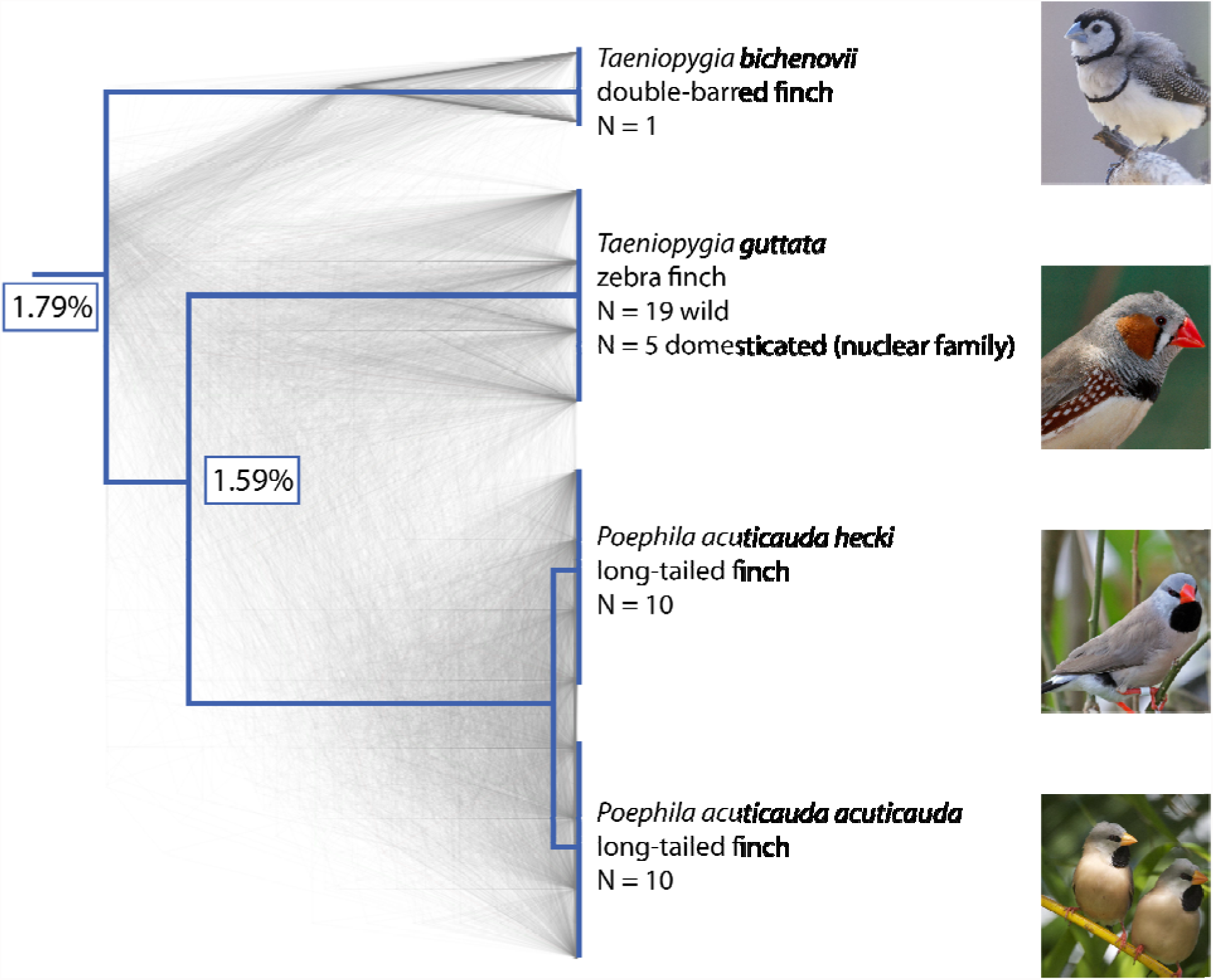
Species tree, gene trees, and sampling numbers for the species in this study: double-barred finch (*Taeniopygia bichenovii*), zebra finch (*T. guttata*), and the two long-tailed finch subspecies (*Poephila acuticauda hecki* and *P. a. acuticauda*). Tree rooted with medium ground finch and collared flycatcher (*Geospiza fortis* and *Ficedula albicollis*; not shown). Shown in gray are 1000 gene trees, which were used to infer the species tree (23). The pairwise divergence between species is indicated at nodes, as measured by the genome-wide average across autosomes. Images of birds from Wikimedia Commons.

We mapped reads from all individuals to the zebra finch reference genome (1 Gb assembled across 34 chromosomes; (*24*)) and generated *de novo* SNP calls for all three species. After filtering for quality, we identified 44.6 million SNPs in zebra finch, 26.2 million SNPs in long-tailed finch, and 3.0 million SNPs in double-barred finch (Table S2). These SNP numbers correspond to autosomal nucleotide diversity of *π*=0.82% and *θ*_w_ =1.37% in zebra finch and *π*=0.55% and *θ*_w_=0.73% in long-tailed finch (*25, 26*), approximately ten times higher than estimates from ape species (*27*). Assuming a mutation rate per base pair per generation of 7 × 10^-10^ *(23)*, this suggests a long-term effective population size (*N*_*e*_) of 4.8 × 10^6^ and 2.5 × 10^6^ in the zebra finch and long-tailed finch, respectively. Thus, these two species have much larger *N*_*e*_ than most other species for which there exist fine-scale recombination maps, with *N*_*e*_ more reflective of biodiversity at large (Fig. S3).

Next, we inferred haplotypes for zebra finch and long-tailed finch using a linkage-disequilibrium approach that incorporated phase-informative reads and family phasing (*28, 29*). From the haplotypes, we estimated fine-scale recombination maps using LDhelmet, an approach that has been shown to perform well for species with higher nucleotide diversity (*14*). The resulting maps estimated genome-wide median recombination rates in zebra finch and in long-tailed finch as ρ = 26.2/kb and 14.0/kb, respectively, corresponding to a median of 0.14 cM/Mb for both species *(23).* Simulations indicated that we had limited power to identify hotspots at either high recombination rates (Fig. S4), so we restricted our analyses to the 18 largest chromosomes in the zebra finch genome (930 Mb; 91% of sequence), ignoring microchromosomes in particular. For these 18 chromosomes, our results accord well with recombination maps inferred from a relatively small pedigree-based study of zebra finch (*30*), with a correlation of 0.91 for rates estimated at the 5 Mb scale (Fig. S5), providing confidence in our inference.

### Hotspots and their evolution

Most of the recombination events in zebra finch and long-tailed finch occur in a narrow portion of the genome, with 82% and 70% of events localized to 20% of the genome in zebra finch and long-tailed finch, respectively (Fig. S6). Given this evidence for heterogeneous recombination, we sought to identify hotspots in the genome. We operationally define hotspots as regions that are at least 2 kb in length, have at least 5-fold the background recombination rate estimated for surrounding 80 kb, and are statistically supported as hotspots by a likelihood ratio test (*31*). This approach yielded 3949 hotspots in zebra finch and 4933 hotspots in long-tailed finch (Fig. 2, S7, S8), with one hotspot detected on average every 215 and 179 kb in the two species, respectively. This difference in hotspot density between species is consistent with simulations that indicate less reliable detection of hotspots when the background population recombination rate is higher, as it is in zebra finch (Fig. S4, Fig. S9). Importantly, the hotspots were detected after aggressively filtering our SNP datasets and show no evidence of having higher phasing error rates than the rest of the genome (Table S4, S5, Fig. S10).

**Figure 2:**
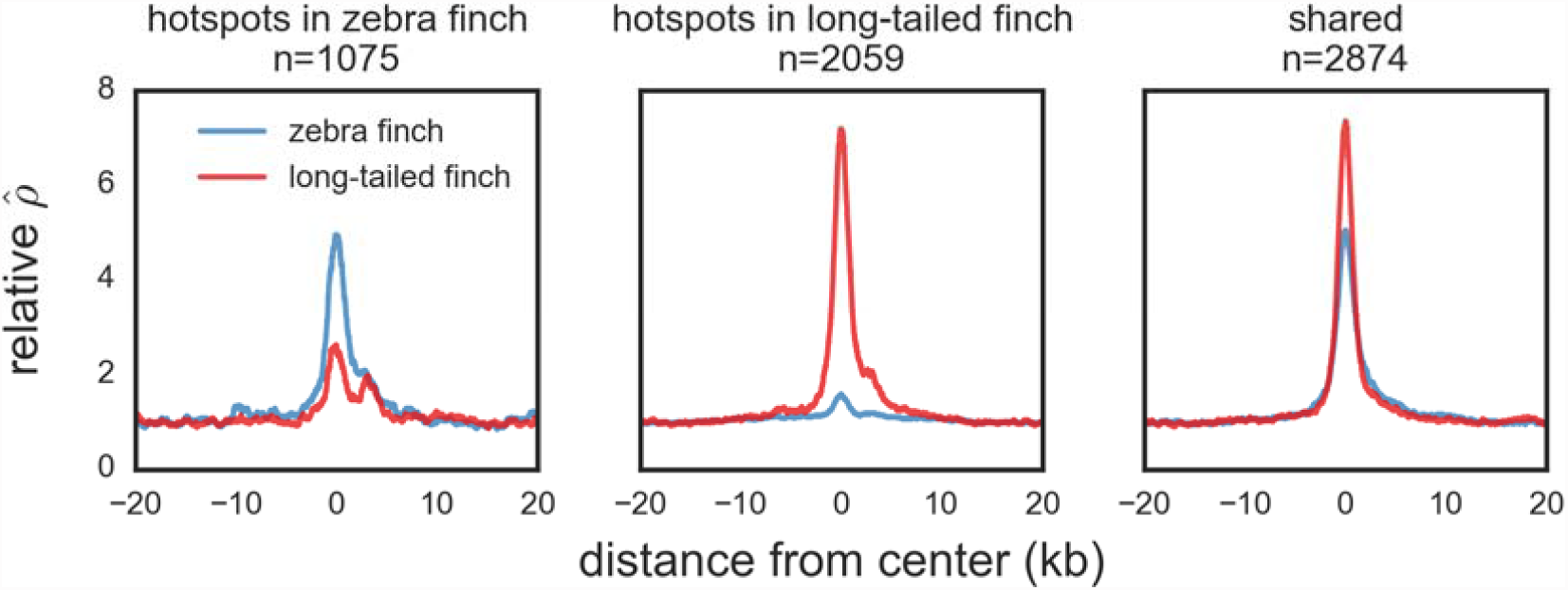
Average relative recombination rate () divided by the background of 20 kb on either side of the hotspot) across hotspots detected only in zebra finch (*Taeniopygia guttata*), those detected only in long-tailed finch (*Poephila acuticauda*), and those inferred as shared in the two species. Shared hotspots are those whose midpoints occur within 3 kb of each other. The orientation of hotspots is with respect to the genomic sequence and has no functional interpretation.

Considering hotspots as shared if their midpoints occurred within 3 kb of each other, 73% of zebra finch hotspots (2874 of 3949 hotspots) were detected as shared between the two species (Fig. S11) when only 3.6% were expected to overlap by chance (Fig. S12); similar results were obtained under different criteria for hotspot sharing (Table S3). The true fraction of shared hotspots between zebra finch and long-tailed finch is likely higher than observed, because we do not have complete power (Fig. S4) and simulations suggest that we are unlikely to detect spurious cases of hotspot sharing *(23)*. On the other hand, the observed levels of sharing are somewhat lower than expected compared to a model in which all hotspots are identical in the two species (Fig. S13).

Such high levels of hotspot sharing contrasts sharply to results from comparative systems in apes and mice, which exhibit no evidence for hotspot sharing even across populations or species with modest levels of genetic differentiation (*9, 10, 32*). As an illustration, when we apply the same criterion as above to humans and chimpanzees, only 10.5% of hotspots overlap when 7.2% are expected by chance (Fig. S12). Interestingly, our findings in birds echo those obtained from four species of *Saccharomyces* yeast, which show nearly complete conservation of hotspot locations and heats across species 15 myr diverged (*33*). Almost all hotspots in *Saccharomyces* yeast occur at promoters, which are evolutionarily stable, suggesting that how hotspot locations are specified influences how they evolve (*34*).

To provide further support for the validity of the inferred hotspots, we tested if they show evidence for GC-biased gene conversion (gBGC), measured as higher equilibrium levels of GC content (GC* (*35, 36*)). Because evidence for gBGC in birds is somewhat indirect (*37, 38*), we first looked for support for gBGC at broad genomic scales, finding a positive relationship between recombination rate and GC* (Fig. 3A-B). Narrowing our focus to the regions surrounding hotspots, we observe that hotspots exhibit peaked GC* relative to both flanking sequence and coldspots matched for the same overall GC and CpG content (Fig. 4A-B). A similar phenomenon is seen in intra-species variation data: at hotspots but not matched coldspots, derived alleles segregate at a higher frequency at AT to GC polymorphisms than at GC to AT polymorphisms (Fig. S14). Thus, two independent signatures of recombination — patterns of linkage disequilibrium and of base composition — converge in demonstrating that finches have recombination hotspots and that these are conserved over much larger time scales than in apes and mice (*8, 9, 39*).

**Figure 3:**
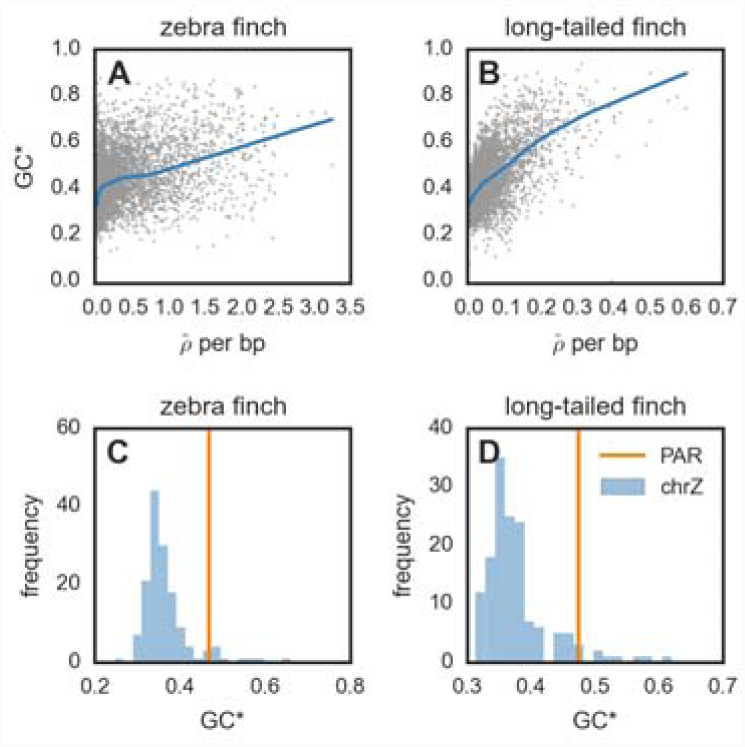
(A-B) Relationship between equilibrium GC content (GC*; 23)) and for zebra finch (*Taeniopygia guttata*) and long-tailed finch (*Poephila acuticauda*) across all autosomal chromosomes. Both GC* and were calculated across 50 kb windows with LOESS curves shown for span of 0.2. (C-D) GC* and the pseudoautosomal region (PAR). The histogram shows GC* for chromosome Z across 500 kb windows; GC* for the 450 kb PAR shown by the vertical line.

**Figure 4:**
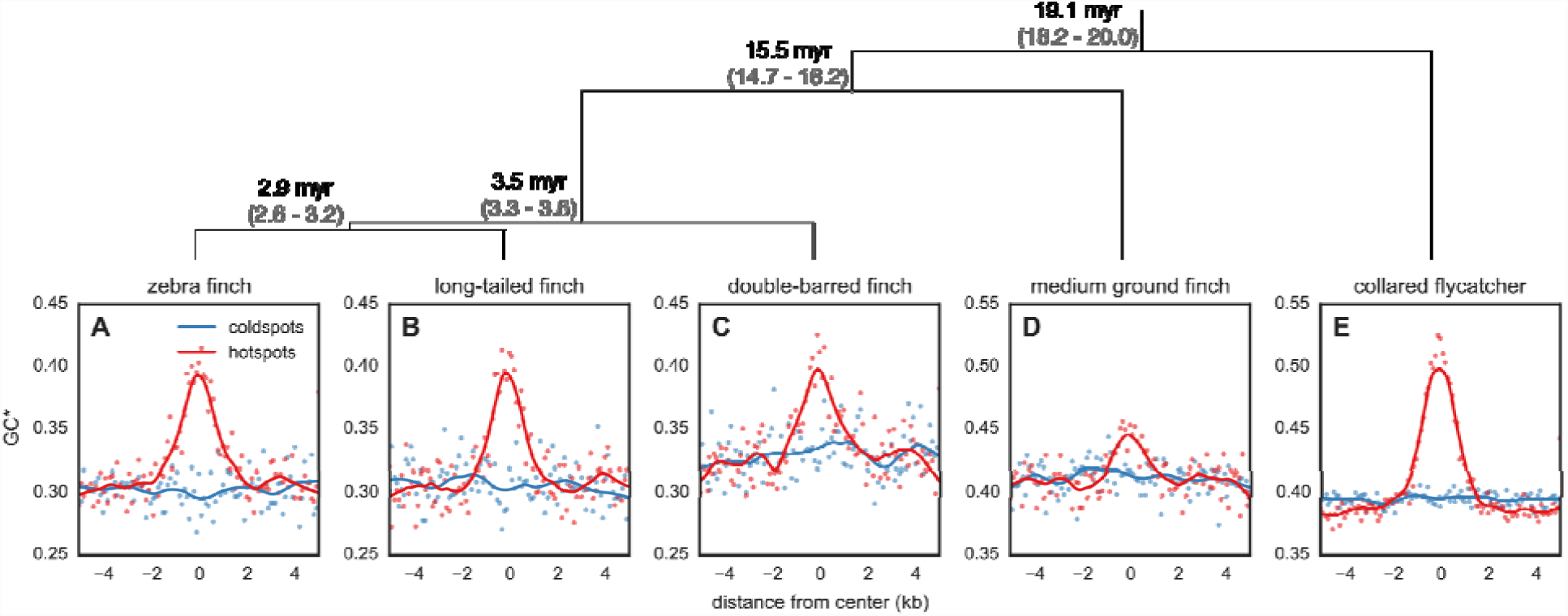
Expected equilibrium GC content (GC*) around hotspots and matched coldspots. Points represent GC* estimated from the lineage-specific substitutions aggregated in 100 bp bins from the center of all hotspots in (A) zebra finch (*Taeniopygia guttata*) and (B) long-tailed finch (*Poephila acuticauda*). GC* for (C) double-barred finch (*T. bichenovii*), (D) medium ground finch (*Geospiza fortis*), and (E) collared flycatcher (*Ficedula albicollis*) was calculated around hotspots identified as shared between zebra finch and long-tailed finch. LOESS curves are shown for a span of 0.2. The orientation of hotspots is with respect to the genomic sequence and has no functional intepretation. Species tree (23) shown with estimated divergence times in millions of years (myr) and its 95% Highest Posterior Density (HPD) in gray.

After observing the pattern of gBGC at hotspots in zebra finch and long-tailed finch, we tested how far conservation of hotspot locations extends across the avian phylogeny by considering three additional species: double-barred finch (an estimated ∼3.5 million years [myr] from zebra finch *(23)*), for which we had collected genomic data, as well as medium ground finch *Geospiza fortis* (∼15.5 myr from zebra finch), and collared flycatcher *Ficedula albicollis* (∼19.1 myr from zebra finch), for which draft assemblies are available (*40, 41*). Because we only had a single diploid genome from these species, we tested for hotspot conservation indirectly by determining if these species had peaks in GC* at the hotspot locations inferred as shared between zebra finch and long-tailed finch. We find localized GC* peaks at hotspots in all three species (Fig. 4C-E), suggesting that the conservation of hotspots extends across tens of millions of years of evolution in birds.

### Analyzing fine-scale rates across the genome

Hotspots in zebra finch and long-tailed finch are enriched near transcription start sites (TSSs), with ∼18% occurring within 3 kb of a TSS (Fig. S15), and near CpG islands (CGIs), with ∼40% of hotspots within 3 kb of a CGI in both species. Consistently, recombination is nearly two-fold higher near TSSs (Fig. 5A-B), and more than three-fold higher near CGIs (Fig. 5C-D). This positive association between proximity to the TSS and recombination rate has been previously reported in a number of species without *PRDM9*, including *S. cerevisiae*, the monkey-flower *Mimulus guttatus*, and *A. thaliana (12, 13, 42)*. In turn, the link between CGIs and recombination rates has been found both in species without *PRDM9* (dogs; (*11*), *A. thaliana;* (*43*)) and, albeit more weakly, in species with *PRDM9* (humans and chimpanzees; (*9*)). Moreover, although the genic recombination rate is positively correlated with expression levels as measured in the testes in the zebra finch (Spearman’s r=0.09, p=2.73×10^-18^, Fig. S16), the relationship between CGIs and recombination rate remains significant after controlling for expression levels (Spearman’s r=-0.11; p=4.32 ×10^-27^). This increase in recombination rates near TSSs and CGIs supports a model in which, particularly in the absence of PRDM9 binding specificity, recombination is concentrated at functional elements that are accessible to the recombination machinery. Indeed, both TSSs and CGIs have been shown to serve as sites of transcription initiation (*44*) and to coincide with destabilization of nearby nucleosome occupancy (*45*).

**Figure 5:**
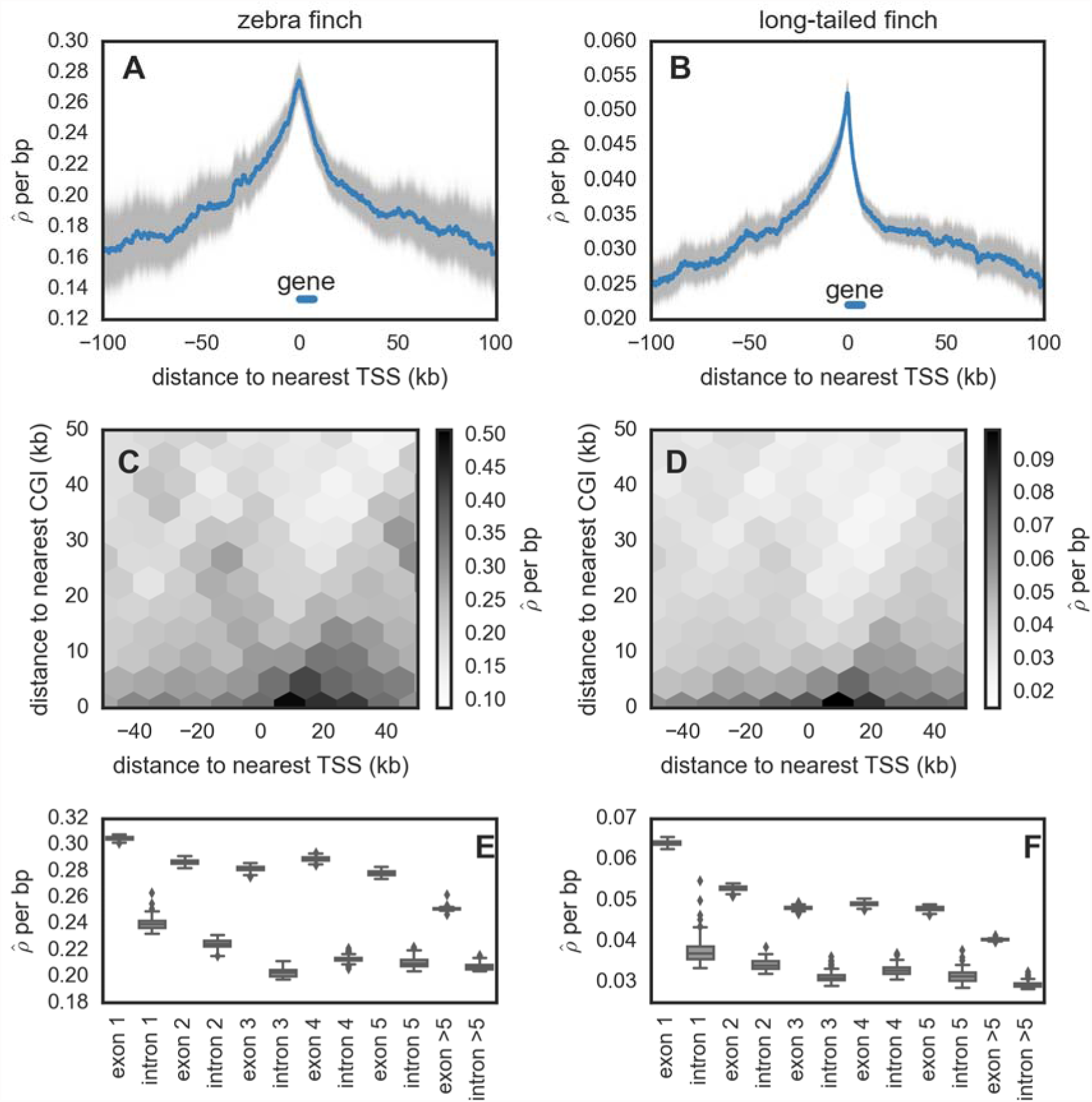
For zebra finch (*Taeniopygia guttata*) and long-tailed finch (*Poephila acuticauda*), (A-B) estimated recombination rates (around transcription start sites (TSSs). The distance to TSS was measured accounting for the 5’ 3’ orientation of genes. Uncertainty in rate estimates (shown in gray) was estimated by drawing 100 bootstrap samples and recalculating means. The median gene length is indicated. (C-D) shown as a function of both distance to the nearest TSS and distance to the nearest CpG island (CGI). Data on CGI in zebra finch were downloaded from the UCSC Genome Browser CGI track and were used for long-tailed finch as well. (E-F) within exons and introns for genes that have 5 exons (n=10,722). See Fig. S29 for simulation results that suggest the inference of higher background in exons does not reflect differences in diversity levels between exons and introns.

Under a model in which the recombination machinery tends to target accessible genomic elements, we would not necessarily expect to see enrichment of specific binding motifs associated with hotspot activity. Accordingly then, when we test for motifs enriched in hotspots relative to coldspots, the top motifs in both species are a string of As, which are also enriched in *A. thaliana* and yeast hotspots and which may be nucleosome depleted or facilitate nucleosome removal (*12, 46*), and a number of additional motifs that appear to be reflective of CGIs (Fig. S17).

At even finer resolution, recombination rates are higher in exonic than intronic regions (Fig. 5E-F), which has also been seen in *A. thaliana* (*13*), dogs (*11*), and monkey-flowers (*42*) and which contrasts to evidence from humans that recombination rates are higher in introns than exons (*47*). One possibility for the pattern seen in the finches is that DSBs preferentially initiate in the first few exons and their resolution occurs in both nearby exons and introns. The specific mechanism by which DSBs would preferentially initiate in exons is unknown, but the pattern is consistent with an important role for chromatin marks that distinguish exons from introns (*45*).

### Contrasting tempos of broad- and fine scale recombination rate evolution

Median recombination rates across and within chromosomes vary over nearly six orders of magnitude (Fig. S9, Fig. S18), creating a heterogeneous landscape of broad-scale recombination rates across the genome, with regions of elevated recombination near telomeres and large intervening deserts (as found in zebra finch pedigree data; (*30*)). This pattern is most pronounced on the sex chromosome Z, which has recombination rates that are more than two orders of magnitude lower compared to the chromosome 1A, the most similarly sized autosome (Fig. S9, (*30*)). While in both zebra finch and long-tailed finch, cytological data indicate that chromosome Z harbors a pericentric inversion polymorphism over most of its length (*48, 49*), such an inversion is unlikely to explain our findings *(23)*.

Between zebra finch and long-tailed finch, broad-scale rates are highly similar, with a genome-wide correlation of 0.82 and 0.84 across 10 kb and 1 Mb windows, respectively (Fig. 6, Fig. S18). Despite this broad-scale concordance, we infer that some genomic regions between the two species have very different rates of recombination (Fig. S19). Moreover, at a greater evolutionary distance, broad-scale patterns differ markedly; unlike zebra finch and long-tailed finch, the collared flycatcher (∼19 myr diverged) has a fairly homogeneous recombination landscape (*50*). This variation in broad-scale rates is particularly notable because, in many species, shifts in broad-scale recombination patterns can be explained almost entirely by chromosomal rearrangements, shifts in karyotypes, and changes in chromosome lengths (*9, 51, 52*). However, there is no obvious pattern by which chromosomal rearrangements drive differences in recombination rates between zebra finch and long-tailed finch (Fig. S19), and, despite harboring a number of small inversions between them, collared flycatcher and zebra finch have similar karyotypes and syntenic genomes (*50*). That broad-scale recombination patterns have changed across the same phylogenetic breadth for which we see hotspot conservation suggests two non-exclusive possibilities: either the heats or locations of some hotspots have evolved, or rates have changed in regions that fall outside of our operational definition of hotspots.

**Figure 6:**
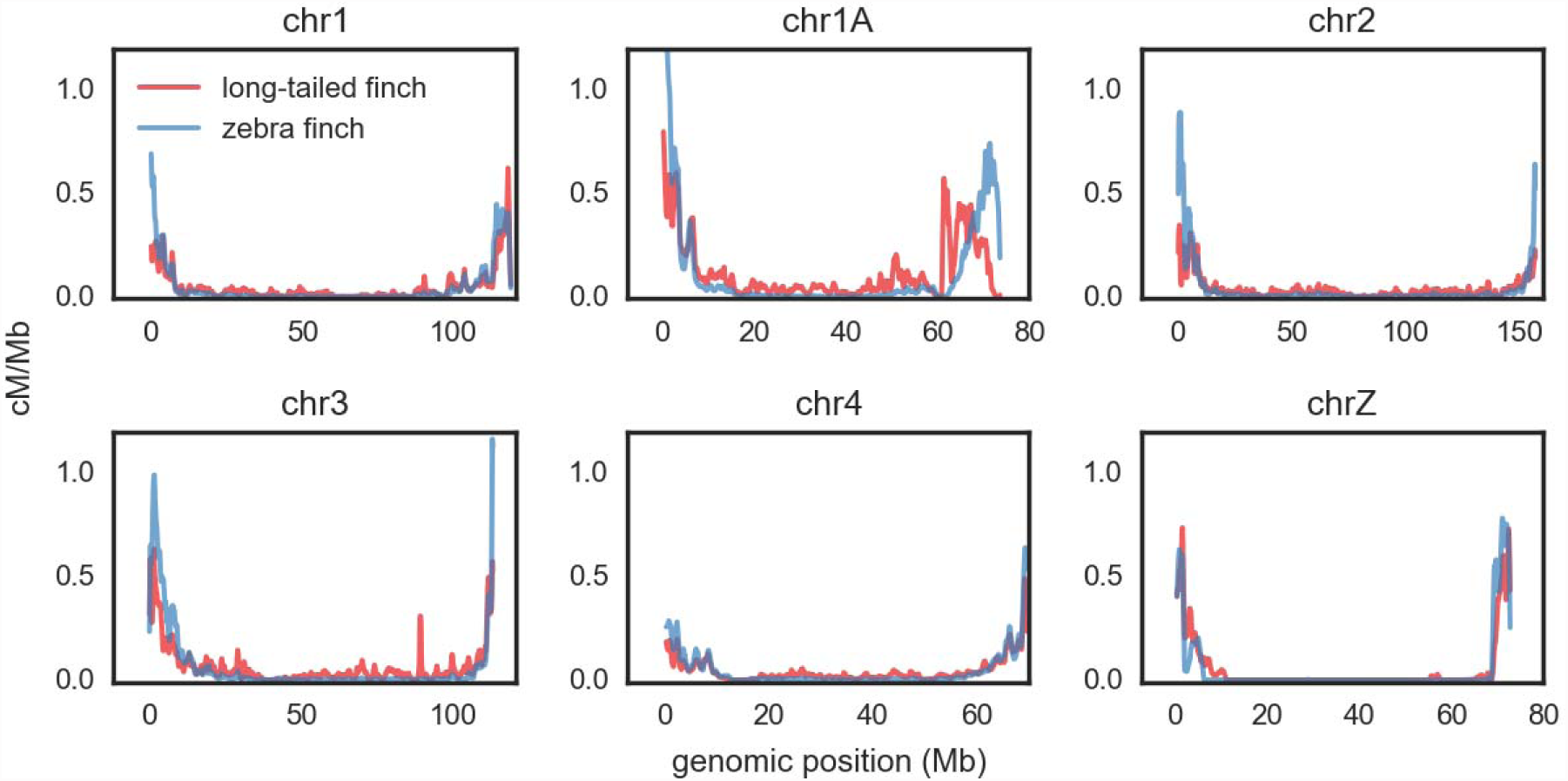
Estimated recombination rate (cM/Mb; obtained from (23)) for zebra finch (*Taeniopygia guttata*) and long-tailed finch (*Poephila acuticauda*), shown as rolling means calculated across 100 kb windows. We show only the five largest autosomal chromosomes and chromosome Z; an expanded version of this figure, showing all chromosomes, is shown in Fig. S18. Rate estimates for chromosome Z should be taken with caution (23).

### Understanding the impact of recombination on the genome

Given the marked variation in recombination rates across the genome, we consider the consequences for genome evolution. First, we note that increased recombination rates drive increased GC content in the genome, presumably via gBGC, and we see this phenomenon both at the genome-wide scale (Fig. 3A-B) and the scale of hotspots (Fig. 4). An extreme example is provided by the pseudoautosomal region (PAR), which we identified on an unassembled scaffold from chromosome Z using estimates of coverage in males and females. We confirmed the PAR by inferring homology to PARs identified in medium ground finch and collared flycatcher (Fig. S20). The PAR is short—estimated to be just 450 kb—and is subject to an obligate crossover in every female meiosis (*53*); as such, it has very high recombination rates. The consequence is visible in the high GC* for the PAR, which exceeds estimates of GC* across most of the rest of chromosome Z in both species (Fig. 3C-D).

Further, as has been reported for many other organisms, notably humans and *Drosophila* (*54, 55*), our results suggest that recombination is positively correlated with levels of nucleotide diversity, particularly on the Z (Fig. S21, S22). This observation is consistent with widespread effects of linked selection in these species (*56*).

## Conclusion

Finches lack PRDM9 yet nonetheless harbor hotspots, with recombination concentrated at promoter features that likely denote greater accessibility to the cellular recombination machinery. In sharp contrast to apes and mice, the hotspot locations are conserved among species several millions of years diverged and likely over tens of millions of years. These results suggest that the genetic architecture of recombination influences the rate at which hotspots evolve. Whereas the binding specificity of PRDM9 drives rapid turnover, the reliance on accessible, functional genomic features leads to stasis. This hypothesis accords with recent results in yeast, in which recombination is concentrated at promoters, and hotspots are stable in intensity and location over tens of millions of years (*33*). To further investigate how deeply this stasis extends and explore the taxonomic generality of these findings, the approaches illustrated here can be applied to the more than forty bird species with available genomes (*19*) and beyond. In doing so, we will begin to better understand why species differ so drastically in their specification of hotspots and, in particular, why a subset rely on PRDM9.

## Acknowledgements

This project was started when MP was a Howard Hughes Medical Institute Early Career Scientist and was funded, in part, by Wellcome Trust grants 086786/Z/08/Z to O.V. and 090532/Z/09/Z to the Wellcome Trust Centre for Human Genetics. We thank Bernard de Massy, Corinne Grey, Simon Myers, Trevor Price, Molly Schumer, Jeff Wall, Amy Williams and John Willis for helpful discussions and/or comments on the manuscript; Karene Argoud and Dr. Paolo Piazza at the Genomics Core at the Wellcome Trust Centre for Human Genetics for assistance with lab work; and M. Thomas Gilbert for sharing the zebra finch gene annotations in advance of publication. We are grateful to Isabel Lam and Scott Keeney for sharing their unpublished results and manuscript with us, and for many helpful discussions.

BAM alignment files for genomic data for zebra finch, long-tailed finch, and double-barred finch are available at XXX. Sequence reads for RNAseq experiments in zebra finch are available at XXX. Filtered variant call files (VCFs) are available for zebra finch, long-tailed finch, and double-barred finch at XXX. Masked genome files for zebra finch, long-tailed finch, and double-barred finch and the reconstructed ancestral genome are available at XXX. Raw recombination output from LDhelmet for zebra finch and long-tailed finch is available at XXX. All scripts and an electronic lab notebook for this work are available at https://github.com/singhal/postdoc and https://github.com/singhal/labnotebook, respectively.

## Supplementary Methods

Materials and Methods

Figures S1-S29

Tables S1-S6

